# Alteration of Soil Bacteriome by Prolonged Exposure to Metal Oxide Nanoparticles

**DOI:** 10.1101/2022.05.16.492223

**Authors:** Nzube Prisca Egboluche, Hongtao Yu, James Wachira

## Abstract

Metal oxide nanoparticles (MONPs) have found applications in many industrial and consumer products and are inevitably released into the environment, including soil. Soils host diverse microorganisms that are integral to ecosystem function including regulating plant growth. In this study, the influence of Cu_2_O, Fe_3_O_4_ and Ag_2_O NPs on soil microbial communities was assessed. Microbial community diversity and compositional structure was characterized using quantitative PCR and 16S rRNA gene sequencing. MONPs altered soil bacteria community composition by causing significant reduction in bacterial diversity and change in bacterial abundance. Soils with Cu_2_O and Ag_2_O NPs treatments significantly reduce bacterial diversity accompanied by shifts at the Class and Phylum taxonomic levels toward bacteria groups responsible for chitin degradation (*Bacteriodetes*) and nitrogen fixation (*alpha-Proteobacteria*). Response of bacterial communities to MONPs exposure is dependent on the exposure time and type of MONPs used. While the mechanisms underlying these observations remain to be elucidated, it is proposed that the known antimicrobial properties of Cu_2_O and Ag_2_O NP_s_ cause reduced growth and viability of some bacteria taxa.

**Importance:** Nanoparticles are finding many applications in society and as such there is the need to gain a better understanding of their potential effects on microorganisms in soil and other environmental niches. Soil contains a large diversity of microorganisms that play many essential roles in organic matter recycling and plant growth. Metagenomics has become an essential tool for understanding the functional diversity of microbiomes and in this study, it was used to assess the diversity of soil bacteria communities in the presence of selected metal oxide nanoparticles. The reported changes in bacterial community structure suggest that nanoparticles have the potential to alter soil microbiomes. However, the effects depend on the composition of the nanoparticles suggesting that environmental impacts of different nanoparticle formulations are necessary.

## Introduction

While nanomaterials arise in natural activities such as volcanic eruptions and the burning of fossil fuels, some are produced for use in diverse fields including food packaging and medical and industrial applications. The impetus to develop nanomaterials has arisen due to some unique characteristics that are not observed in similarly formulated bulk materials such as new optical, electrical, and chemical properties^3^. Whereas the scientific basis for these properties remain to fully understood, the size and the surface area of nanoparticles (NPs) are critical factors that influence the reactivity of NPs ^2,4^. NPs are used in different applications because of their desired properties including optical^5^, mechanical^6^, magnetic and chemical^1^. NPs have been found useful in many fields of application including medical, pharmaceutical, electronic, food and cosmetics industries^1,7–10^. With increased production and use of different types of NPs, such as metal and metal-oxide NPs, it has resulted in continuous disposal of these NPs into the environment. Disposed NPs mostly end up in the soil and their accumulation could lead to exposure to living organisms^11^, thus raising concerns on the potential contamination of the environment and thus increased ecological risk.

Microbes are ubiquitous in the environment where they play diverse roles that reflect on both natural and perturbed functioning of ecological niches. Soil is a complex medium that hosts diversity of microorganisms, bacteria, fungi and archaea. The complexity, functioning, and process**es** of the soil ecosystem is driven by soil microorganism and their extracellular enzyme activities^12^. The microbial diversity of soil microorganisms is one of the major indicator parameters of soil quality and productivity^13^. Soil microbiomes are influenced by environmental stressors resulting in changes in soil physical and chemical properties, thus altering the ecosystem functioning. The accumulation of heavy metals (Cd, Cu, Pb, & Zn) in soil led to overall microbial diversity decrease^14^. Given the substantial increase in the application of NPs in commercial products such as cosmetics, coatings, paints, food additives, drugs and electronics, release into the environment becomes inevitable.

The release of NPs into agricultural soils could be either deliberately as agrochemicals (e.g., nanopesticides)^15^ or accidentally as contaminants^16,17^. The incorporation of NPs such as C_e_O_2_, Ag, ZnO, Cu_2_O in agrochemicals including fertilizers, pesticides^10,15,18^ or growth regulators^19^ have been reported. Many metal and metal oxide NPs, such as Ag, Cu_2_O, TiO_2_ and ZnO NPs, have intrinsic antimicrobial properties, and therefore could be toxic to beneficial bacterial and fungi^11,19,21^. For example, soil bacterial communities decreased at day 15 and 60 in TiO_2_ and ZnO NPs treated soils but had no effect at day zero^22^ suggesting the impact of long-term exposure of NPs to microbial abundance. Others have reported significant reduction in bacterial and archaeal abundance with Ag NPs while Al_2_O_3_ and SiO_2_ NPs had no effect on microbial composition and community^23^. Alterations in microbial community structure and functioning by NPs have also been reported^24,25,26^. The *Actinobacteria* and *Acidobacteria* groups have been found to be susceptible to AgNPs application while *Bacteriodetes* and *Proteobacteria* were resistant^23,27^. In a planted and unplanted soils, AgNP at 100 mg/kg soil altered the structural composition of bacterial community and the soil metabolites^28^. In a study in which AgNP were added to soil in a mixture with sewage slurry, one species of plants (*Microstegium vimineum*) was adversely affected as was the microbial biomass^29^. The soil microbial ecosystem also exhibited changes in species composition as measured with operational taxonomic units (OTU_s_). Interestingly, these authors reported that the NPs caused higher magnitudes of change than a AgNO_3_ positive control.

Considering the broad application and continuous release of NPs in the environment, there is still sparse knowledge on the potential impact to soil microbial communities. While soil microbial communities are significant in maintaining ecosystem functioning such as plant growth, this study sought to investigate the response of soil microbial communities to the presence of different MONPs formulations. Thirty days incubation study was conducted using garden soil and Cu_2_O, Ag_2_O, and Fe_3_O_4_ NPs at 500 mg/kg. The effect of MONPs on known soil bacterial phyla alpha- and *beta-Proteobacteria, Bacteriodetes, Actinomycetes*, and *Firmicutes*, was quantified using quantitative real-time PCR (qPCR) while the overall biodiversity was characterized using 16S rRNA gene sequencing.

## Materials and Methods

The copper oxide (Cu_2_O), silver oxide (Ag_2_O) and iron (II, III) oxide (Fe_3_O_4_) NPs in powder form were used in this study. The Cu_2_O and Fe_3_O_4_ NPs were purchased from the U.S. Research Nanomaterials, Inc. The characterization of these NPs was adopted from the manufacturers. Cu_2_O NP has a diameter size of 10 nm, spherical shape, brown-black color, and 99% purity, while Fe_3_O_4_ NPs had a diameter size of 15-20 nm, dark brown color, spherical shape and 99.5% purity. Zeta potential is about 10.8 mv for pH 7. The Ag_2_O NPs in powder form was obtained from Sky Spring Nanomaterials Inc. Based on the manufacturer’s characterization report, Ag_2_O NPs had a diameter of 20-30 nm, spherical shape, gray color and 99.9% purity.

### Soil Incubation and Microbial DNA Extraction

Garden soil was thoroughly mixed with Cu_2_O, Ag_2_O, and Fe_3_O_4_ NPs at 500 mg/kg respectively in 5 replicates for each treatment, then placed in a controlled environment between 25°c - 30 °C to incubate. At day 1 of the incubation (24 hours after MONPs addition) and at day 30, soil samples were collected from soil-NP treatments and control (0 mg/kg). Soil microbial DNA was extracted from the soil samples using DNeasy PowerMax soil isolation kit (MOBIO Laboratories, CA, USA), following the manufacturer’s instructions. The concentration of the extracted DNA was determined with Qubit 4 fluorometer that uses double-stranded DNA binding dye.

### Quantitative Real-Time Polymerase Chain Reaction (qPCR)

Real-time qPCR was performed to quantify the known bacterial phyla, alpha- and beta-*Proteobacteria, Bacteriodetes, Actinomycetes*, and *Firmicutes* using bacteria phylum specific primers (Table 1)^30^. SYBR Green master mix fluorescent dye that is highly sensitive to DNA and phylum specific bacteria primers including the DNA template were used for the qPCR. The mock microbial DNA standard obtained from Zymo research was used for optimization while a no template control was used as a negative control. DNA samples at a volume of 1 μL, forward and reverse primers at 2.5 μL respectively, SYBR Green as master mix at 10 μL and nucleases free water at 4 μL were used to perform 20 μL PCR reaction at standard setting. The amplification condition was as follows: denaturation at 95 °C for 60 s, annealing temperature was based on the melting temperature of the specific primers for 60 °C and extension 60 °C for 60 s at 40 cycles. Finally, a melting curve analysis was performed from 60 °C to 95 °C with a ramp rate of 0.1^0^C. The qPCR data containing the melt curve and threshold cycles were exported. One-way ANOVA analysis with a Tukey’s HSD test to compare sample means was conducted. Probability level p < 0.05 was considered statistically significant. The figures were created using the GraphPad Prism 6 software (GraphPad Software Inc.).

**Table 1.**
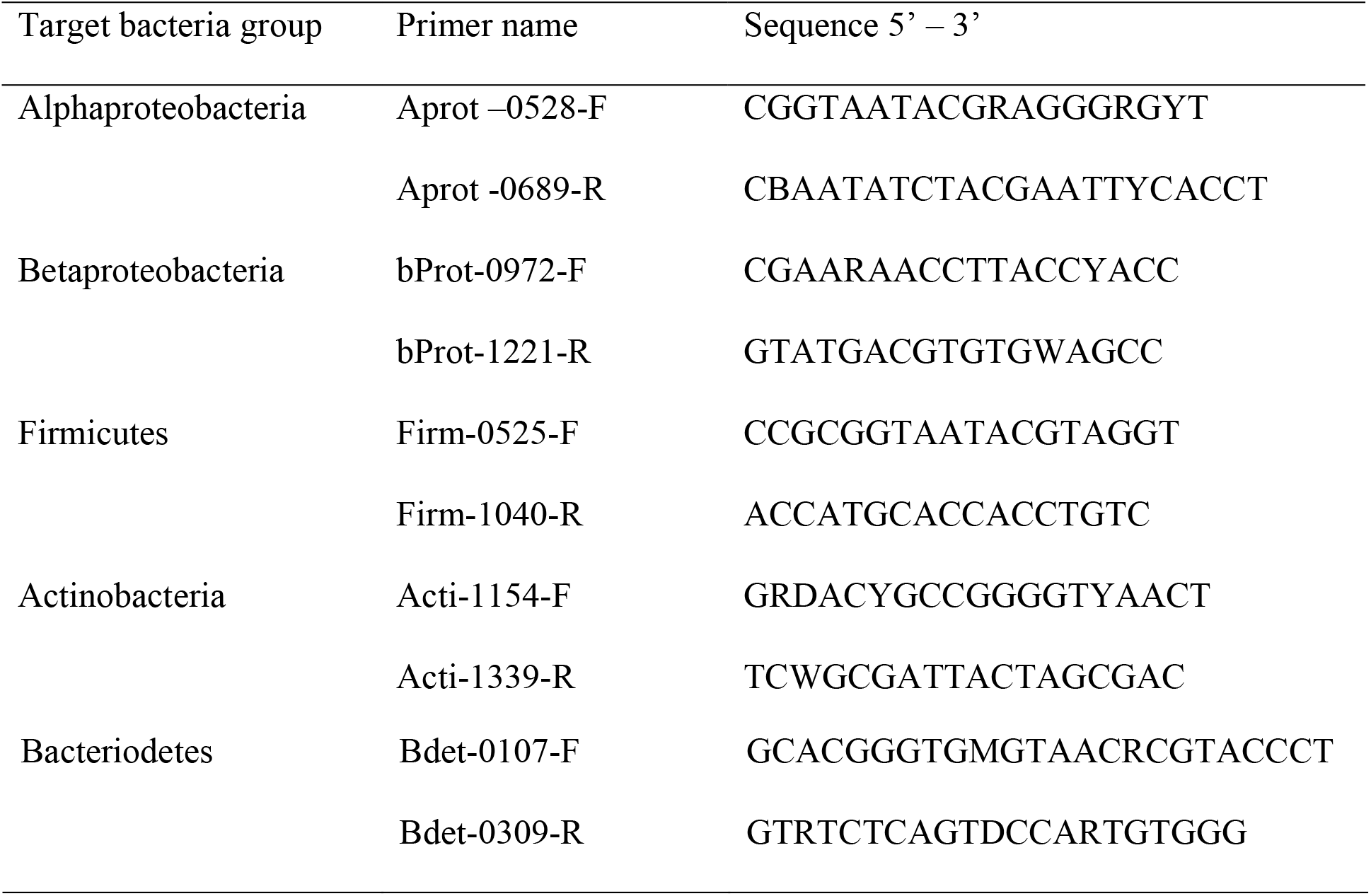
The list of bacteria phylum specific primers that targets some common bacteria groups found in agricultural soils.

### Illumina-Based 16S rRNA Gene Sequencing

Soil microbial community diversity and composition was characterized using high throughput 16S rRNA sequencing on illumina MiSeq instrument. The concentrations, size and quality of the stored DNA isolates were determined using Qubit 4 fluorometer and tape station respectively. The V3 – V4 regions of the bacterial 16S rRNA gene were amplified with 1 μM of 16S primers 5’-TCGTCGGCAGCGTCAGATGTGTATAAGAGACAGCCTACGGGNGGCW-GCAG-3’ forward, 5’-GTCTCGTGGGCTCGGAGATGTGTATAAGAGACAGGACTACHVG-GGTTCTAATCC-3’ reverse and 2x KAPA HiFi HotStart master mix using a thermocycler PCR system (GeneAmp 9700, ABI) following the illumina 16S metagenomic sequencing library preparation protocol. The library product was purified, and final product size validated using a tape station. The libraries were quantified using Qubit 4 fluorometer, normalized and pooled in equimolar amounts. The amplicon sequence was done as a paired-end sequence (2 x 300) on a MiSeq instrument (Illumina, San Diego) according to the illumina standard protocol. Quality control of the data was also managed with FastQC^31^, followed by summarization with MultiQC^32^. Based on the quality metrics, the sequences were filtered and trimmed for further analysis. The sequences were demultiplexed and analyzed using Mothur (version 1.39.1) program following an established workflow developed by Schloss^33^. Using SILVA 16S rRNA database as a reference gene database, sequence reads were blasted for identification and classification. Related sequences were clustered and the number of representatives of each cluster counted which is referred to as operational taxonomic units (OTUs) at 97% sequence similarity. The OTU ID were assigned to different taxonomic levels which produced a taxonomy table that was used for downstream analysis and visualization.

### Soil Microbial Community Diversity Measurement

Using the taxonomy table, microbial diversity measurement was conducted. This involves estimating the alpha and beta diversity of the microbial communities based on the weighted UniFrac pairwise distance matrix of all samples at two-time points day 1 and 30. The alpha diversity was estimated using the Chao1 et al^34^ and Simpson index^35^. To derive the dissimilarity of microbial communities (beta diversity) from different sample treatments the non-metric multi-dimensional scaling (NMDS) and principal coordinate analysis (PC_o_A) metric multi-dimensional scaling analyses were conducted. The Bray-Curtis Emperor plot was also generated with QIIME2 plugins including the DADA2 denoising pipeline^36,37^. The statistical significance of the microbial diversity was compared using analysis of molecular variance (AMOVA) and homogeneity of molecular variance (HOMOVA) statistical analyses. Both AMOVA and HOMOVA analyses were done in Mothur. The visualization of different phylum and class of bacteria composition and phylogenetic tree construction were done in R using phyloseq workflow developed by McMurdie and Holmes^38^ and web-based galaxy software.

## Results

### Effects of MONPs on Soil Bacteria Phyla

Real time quantitative PCR (qPCR) and high throughput 16S rRNA gene sequencing were used to assess the impact of MONPs on soil microbial communities. Five bacteria phyla (Table 1) commonly found in agricultural soil were quantified with qPCR. After 30 days of incubation, the qPCR result showed that *Proteobacteria* was dominant in all the soil samples irrespective of the MONPs treatment. On day 1, there was no significant difference in the bacterial abundance between MONPs and control treatments (Fig. 1a) while on day 30, Cu_2_ONP treatment significantly altered the abundance of the target bacteria groups particularly the *Actinomycetes* and *Bacteriodetes* when compared to control (p<0.02) and Fe_3_O_4_NP treatments (p<0.003) (Fig. 1b). Among the MONPs treatments, Fe_3_O_4_NPs had lower Ct value compared to other MONPs treatment types, indicating that the addition of Fe_3_O_4_NPs increased the overall abundance of the target bacteria groups. The addition of Ag_2_ONPs and Cu_2_ONPs increased the abundance of *Bacteriodetes* when compared to control (Fig. 1c). Overall, soil bacterial groups responded differently to the different MONPs treatments.

**Figure 1.**
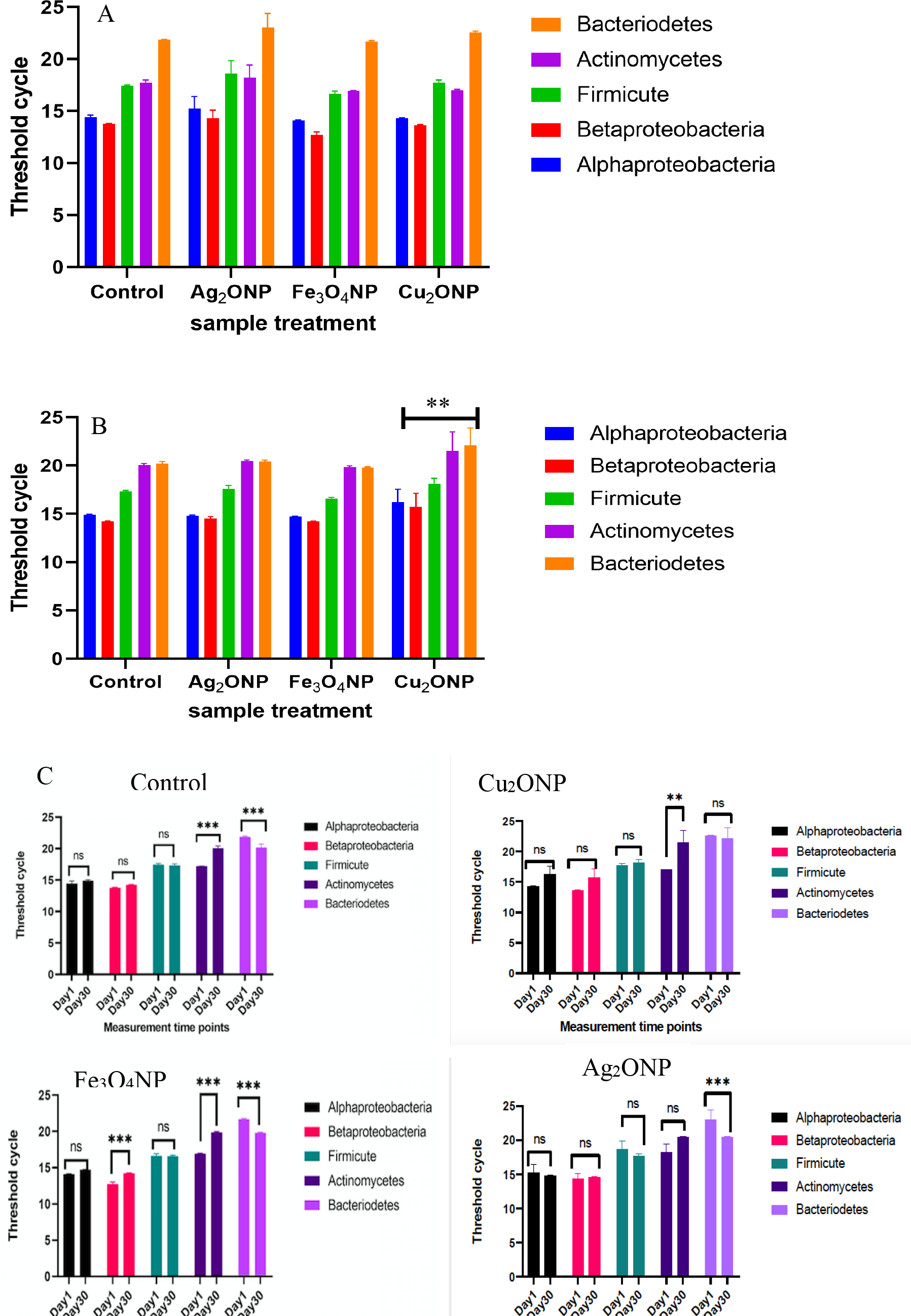
The abundance of target bacteria phyla in the soil exposed to Cu_2_O, Fe_3_O_4_, and Ag_2_O NPs at 500 mg/kg and control (0 mg/kg) measured by their threshold cycle on the y-axis and sample treatments on the x-axis at (A) day 1 & (B) day 30. C) Relative abundance of target bacteria in the soil measured by their threshold cycle on the y-axis and time period on the x-axis in different soil treatments. Each bar represents the mean (n=5) and the standard error within each sample treatment. The lower the threshold value the higher the proportion of the bacteria group.

**Table 2.** The ANOVA p-values of the target bacteria and the time of measurement - day 1 & 30 under each treatment.

| Source of variation | Control | Ag<sub>2</sub>ONP | Fe<sub>3</sub>O<sub>4</sub>NP | Cu<sub>2</sub>ONP |
| --- | --- | --- | --- | --- |
| Time period | <0.0001<sup>(***)</sup> | 0.5609<sup>ns</sup> | <0.0001<sup>(***)</sup> | 0.0164<sup>(*)</sup> |
| Target bacteria | <0.0001<sup>(***)</sup> | <0.0001<sup>(***)</sup> | <0.0001<sup>(***)</sup> | <0.0001<sup>(***)</sup> |
| Interaction<br>(time x target bacteria) | <0.0001<sup>(***)</sup> | 0.0832<sup>ns</sup> | <0.0001<sup>(***)</sup> | 0.1895<sup>ns</sup> |

ANOVA significant at p<0.05, (ns) means not significant.

### Metagenomics Analysis of Microbial Composition

The 16S rRNA sequencing result showed that a total of 15,170,036 high-quality sequence reads were obtained from all the treatments. The sequence reads ranged from 1,323,255 – 2,696,830 among the treatment groups. The bacteriome was dominated by *Proteobacteria, Bacteriodetes, Acidobacteria, Chloroflexi, Actinobacteria, Planctomycetes, Chlorobi, Verrucomicrobia, Firmicutes* and other unclassified bacteria group at varying relative abundance at different time points (Table 3). *Proteobacteria, Bacteriodetes*, and unclassified bacteria groups made up of 76% of the total bacteria population (Table 3; Fig. 2). Similar to the qPCR result with the 16S analysis, *Proteobacteria* was found to dominate the soil samples consisting approximately 40% of total bacteria phyla. The class of *Proteobacteria* comprised of *alpha-Proteobacteria, gamma-Proteobacteria* and least abundance of *beta-Proteobacteria* (Fig. 3). Phylum *Parcubacteria* (OD1) was only present in control treatment indicating decreased abundance by MONP addition. There was no change in the abundance of *Firmicutes* at different time points (Table 3).

**Table 3.** Top 10 bacteria phyla present in the soil and their percentage abundance at day 1 and day 30.

| Bacteria Phyla | Day 1 | Day 30 |
| --- | --- | --- |
| Proteobacteria | 39.5% | 37.8% |
| Bacteria-unclassified | 24.8% | 24.9% |
| Bacteriodetes | 11.3% | 13.9% |
| Acidobacteria | 6.7% | 6.9% |
| Chloroflexi | 3.8% | 3.1% |
| Actinobacteria | 3.6% | 3.4% |
| Planctomycetes | 2.2% | 1.7% |
| Chlorobi | 2.1% | 1.4% |
| Verrucomicrobia | 1.5% | 1.8% |
| Firmicutes | 0.9% | 1.0% |

**Figure 2.**
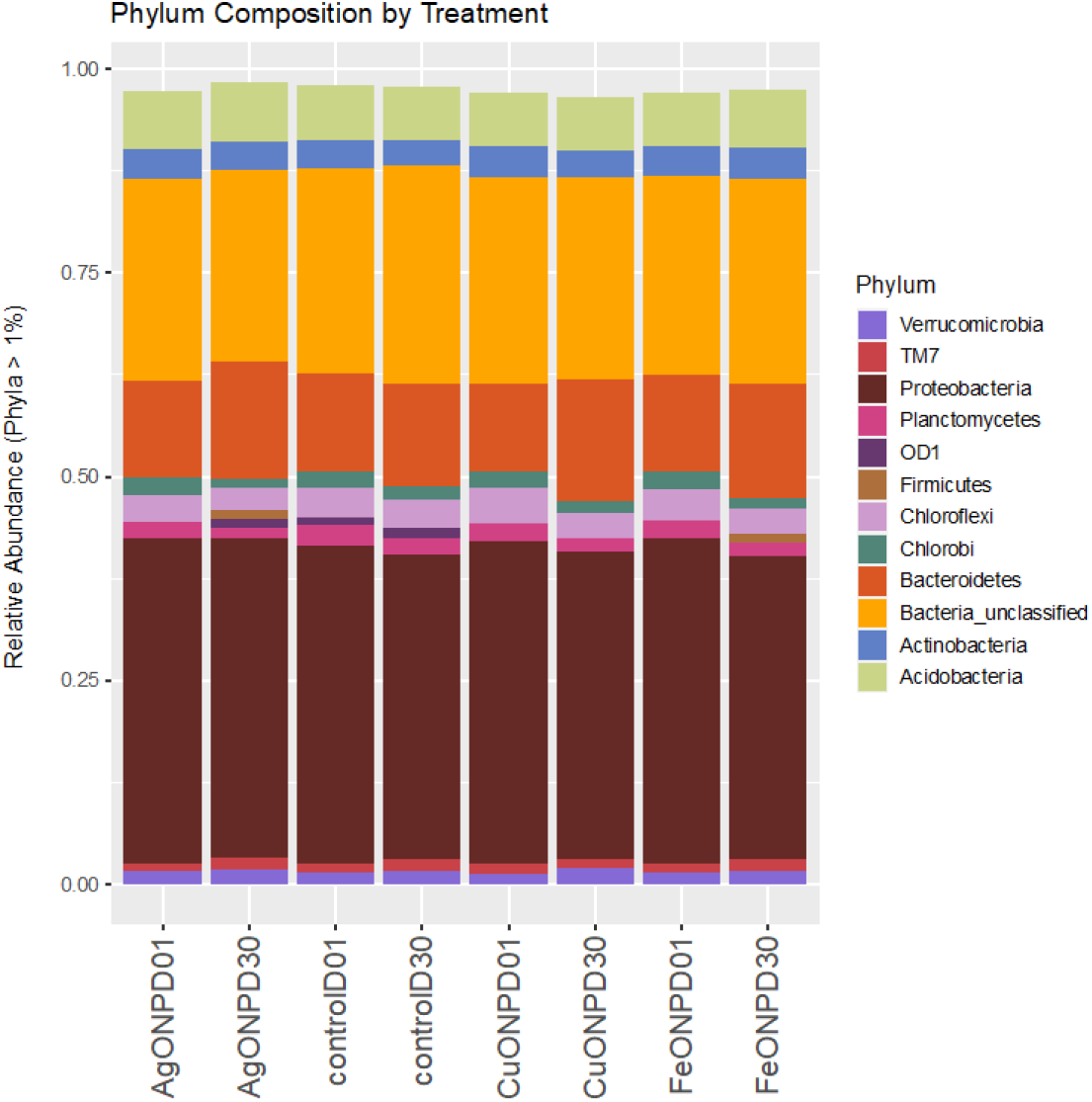
Relative abundance of different bacteria phylum in soil treated with Cu_2_O, Fe_3_O_4_, and Ag_2_O NPs and control (untreated soil) at day 1 and day 30.

**Figure 3.**
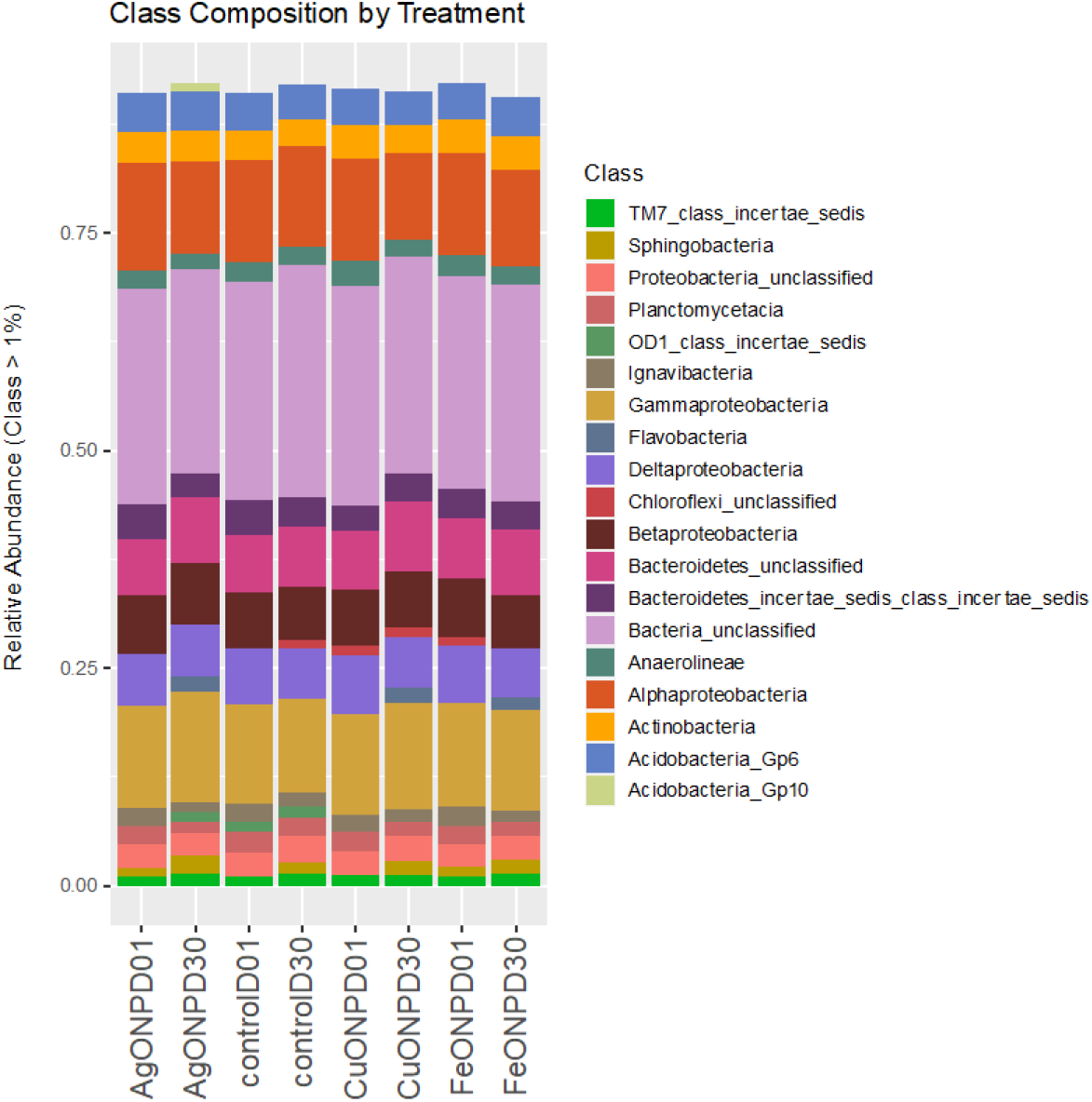
Relative abundance of different classes of bacteria in soil treated with Cu_2_O, Fe_3_O_4_, and Ag_2_O NPs and control (untreated soil) at day 1 and day 30.

Sequence reads ranged from 1,763,083 to 2,696,830 on day 1 and day 30 respectively in untreated soil samples while in MONPs treated soils, the number of reads decreased ranging from 1,323,255 to 1,943,646. On day 1, the total number of OTUs richness was 50,395 and 41,995 on day 30 (Fig. 4 a & b). Compared to control treatments, bacterial OTUs significantly decreased with the addition of MONPs (Fig. 4, Table 4). The population of *Bacteriodetes* had a 2 - 4% increase after 30 days of incubation in all the MONP treatments but remained unchanged in the control. While the relative abundance of *Chlorobi* and *Parcubacteria* (OD1) phyla were negatively affected by MONPs, there was no change in the abundance of *Firmicutes* across all the treatments at different time points (Table 3). Different classes of bacteria were also affected by the MONPs like the *Acdiobacteria-Gp10, Chloroflexi-unclassified, OD1 class_incertae_sedis* and *Flavobacteria* (Fig.3).

**Table 4.** Analysis of molecular variance (AMOVA) and Homogeneity of molecular variance (HOMOVA) of microbial communities.

| Analysis | Comparison | P-value |
| --- | --- | --- |
| AMOVA | Day 1 – Day 30 | 0.031* |
|  | Ag<sub>2</sub>ONP-Cu<sub>2</sub>ONP-Fe<sub>3</sub>O<sub>4</sub> NP-Control | 0.796<sup>ns</sup> |
| HOMOVA | Day 1 – Day 30 | 0.035* |
|  | Ag<sub>2</sub>ONP-Cu<sub>2</sub>ONP-Fe<sub>3</sub>O<sub>4</sub> NP-Control | <0.001* |

**Figure 4.**
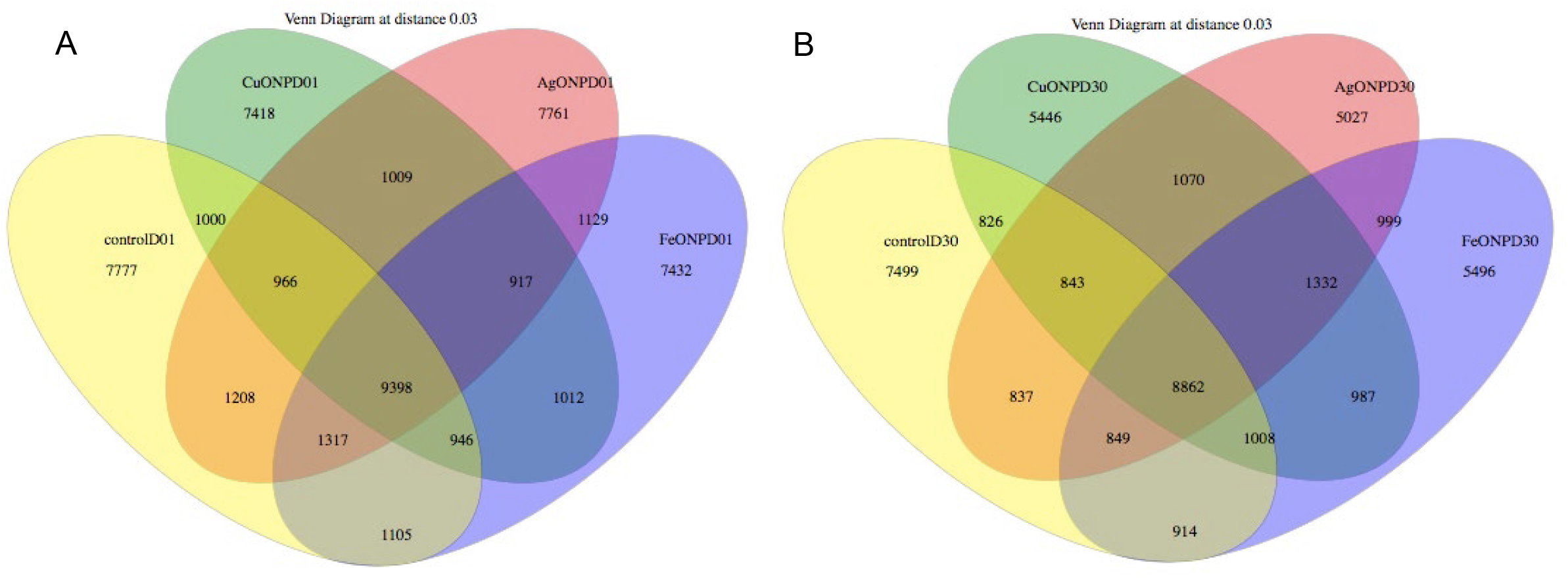
Venn diagram showing the amount of individual and shared OTUs within, between and among 4 sample treatments Cu_2_O, Fe_3_O_4_, and Ag_2_O NPs and control (untreated) on a) day 1 and b) day 30. The number of OTU richness decreased on day 30 for the MONPs treatments.

### Soil Bacteria Community Diversity Change with MONPs Exposure

The alpha diversity of bacteria community within the sample treatments was measured using the Chao1 and Simpson index metrics (Fig. 5). The Chao1 diversity index showed that in the control treatment, the bacteria communities were intact and stable after 30 days there was no significant reduction in bacteria richness. For the MONPs treated soils, the Chao1 diversity index value decreased over time with Ag_2_ONP having the lowest Chao1 index value on day 30 and Cu_2_ONP on day 1 (Fig. 5). The principal coordinate analysis (PC_o_A, Fig. 6a) and non-metric multi-dimensional scaling (NMDS, Fig. 6b) showed the difference in microbial diversity between sample treatments at day 1 and day 30 (p < 0.031, AMOVA analysis). On day 1, the MONP treatments and control samples were clustered together as shown in Fig. 6a. On day 30, the MONPs treatments were significantly distant from the control indicating a significant change in the bacteria community diversity with the addition of Cu_2_O, Fe_3_O_4_, and Ag_2_O NPs (p < 0.0001, HOMOVA analysis) (Table 4). The hierarchical clustering of the treatments also showed that on day 30, control sample was distant apart from the MONPs treatments (Fig. 6c). Of note is that the Bray-Curtis dissimilarity index clusters the samples based on time indicating that the microbial composition changed as a function of time irrespective of treatment (Fig. 6d). This observation is reflected in diversity distance measurement using weighted UniFrac (Fig. 5), which demonstrates the community differences between the two time-points for most of the variation in the data (Fig. 6).

**Figure 5.**
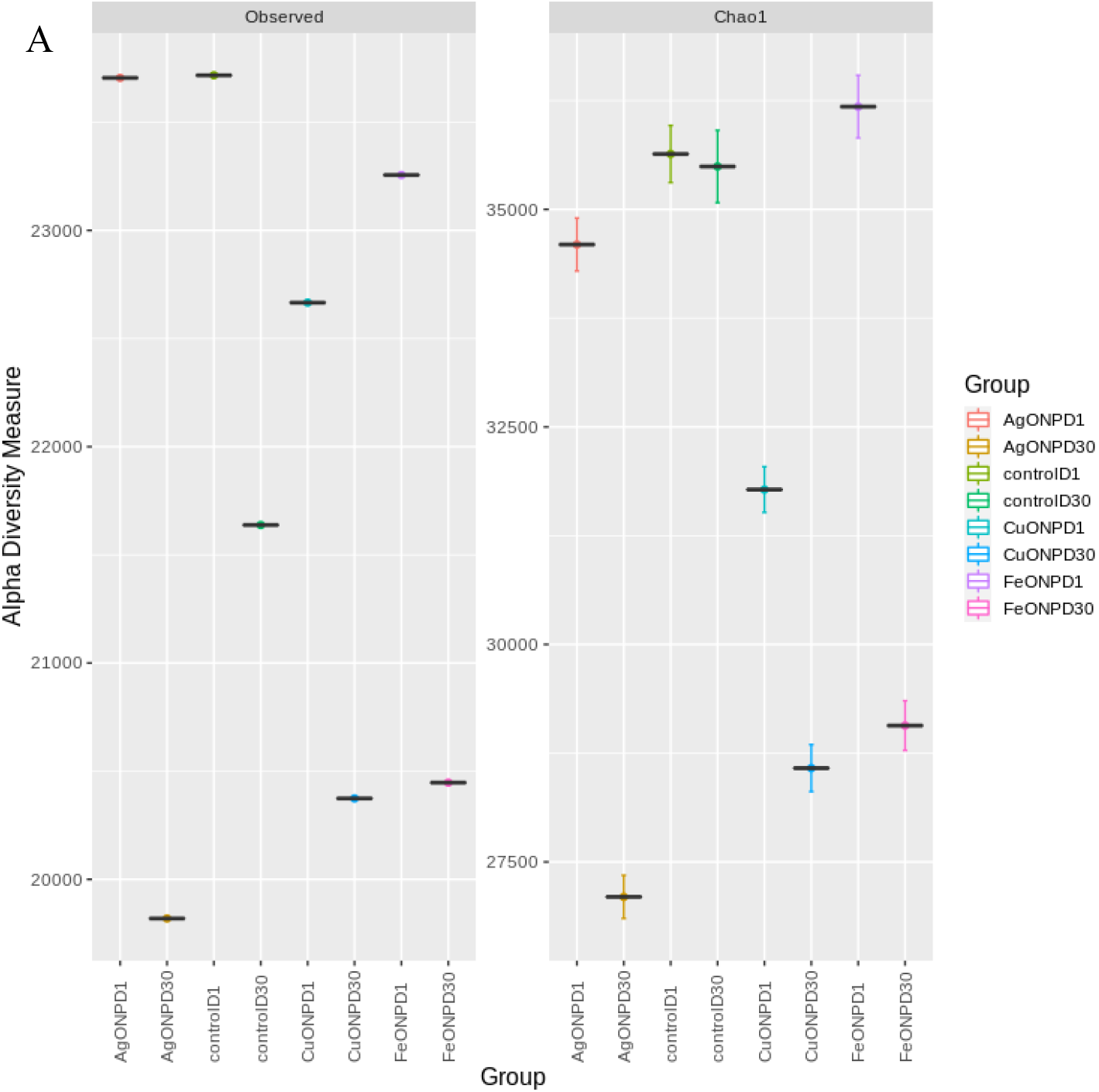
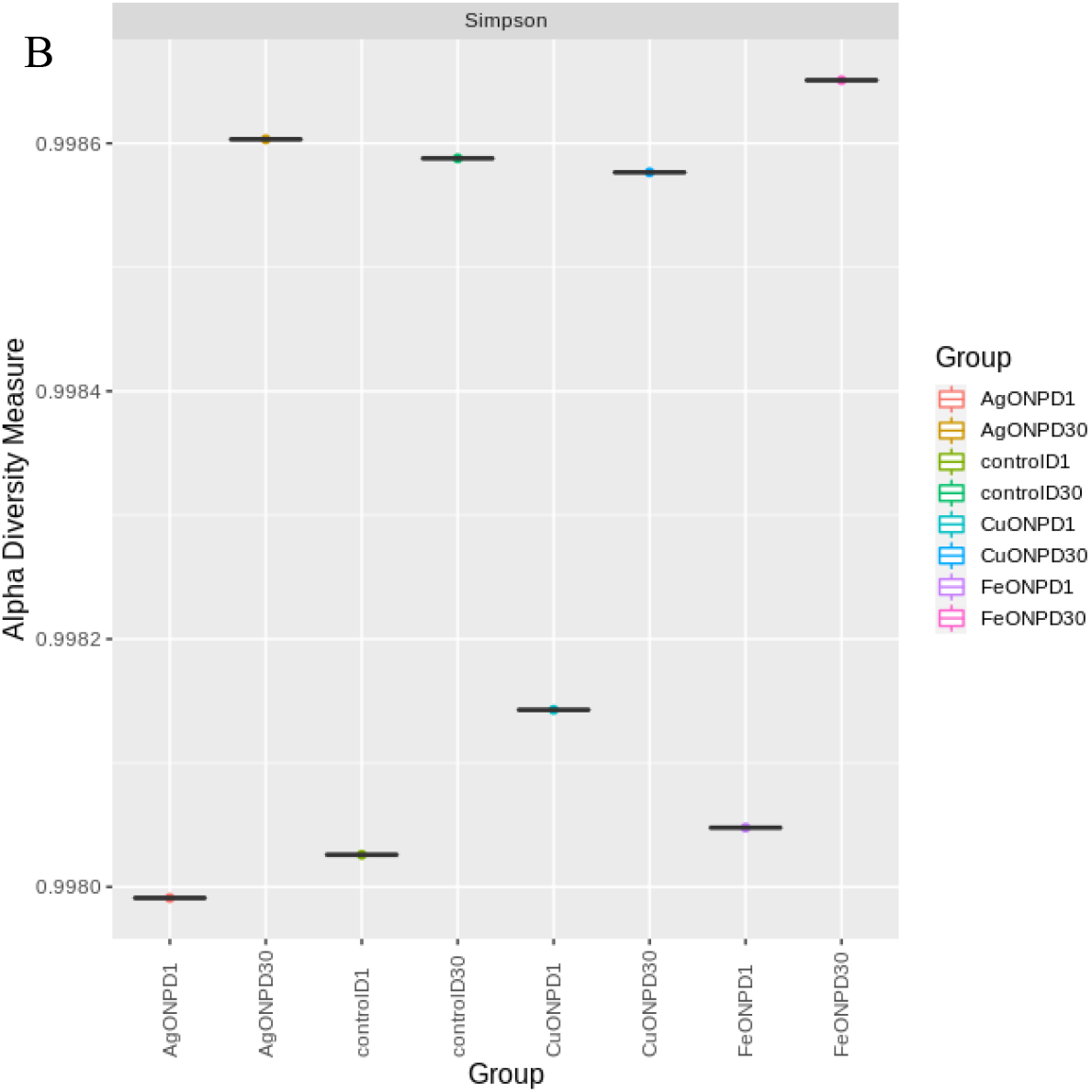
Alpha diversity measurement by a) Observed and Chao1 indices and b) Simpson index to estimate the bacteria community richness and evenness respectively after 30 days of soil exposure to Cu_2_O, Fe_3_O_4_, and Ag_2_O NPs. Bacteria community richness were measured on day 1 and day 30 within each sample treatment.

**Figure 6.**
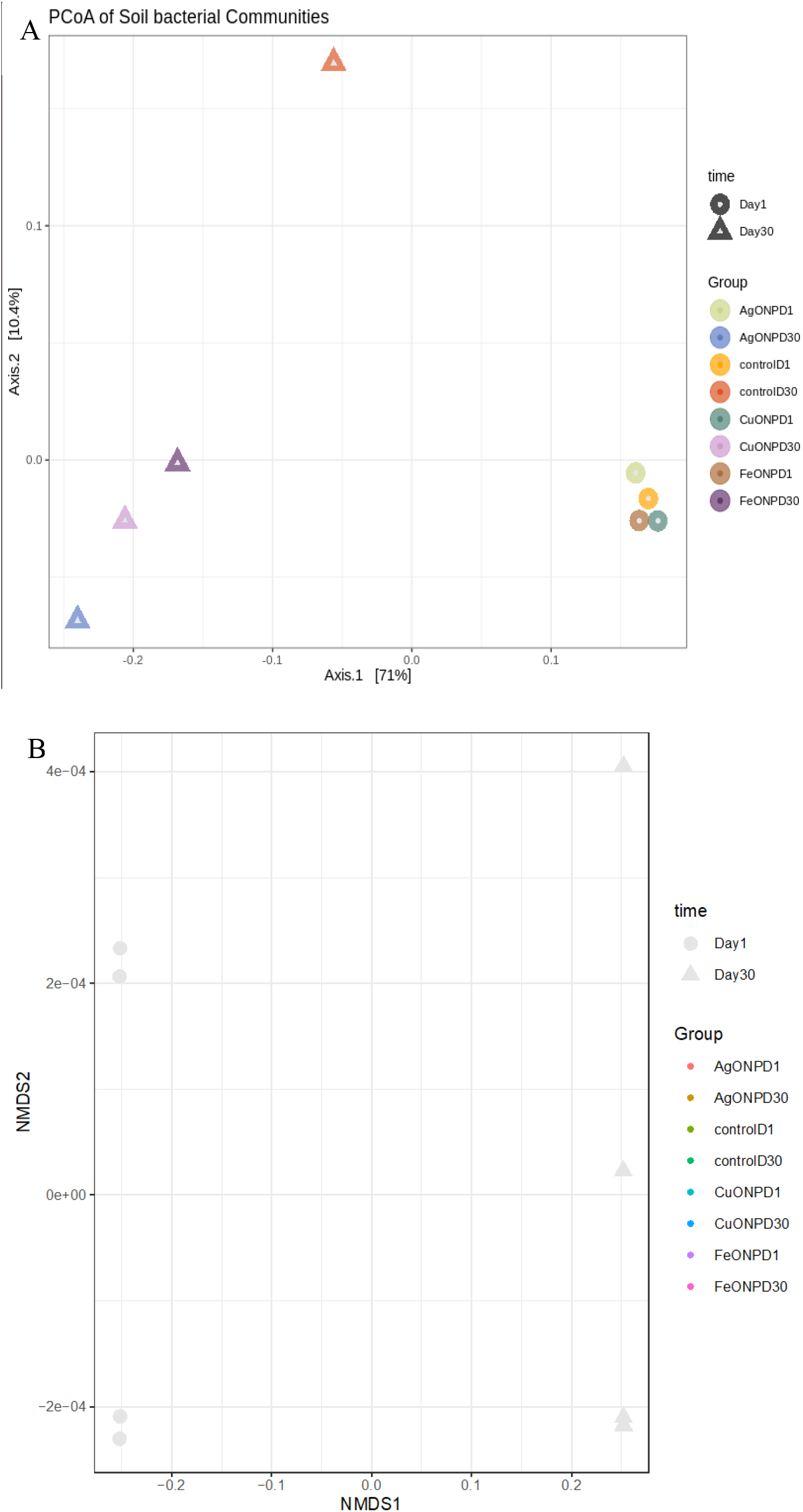
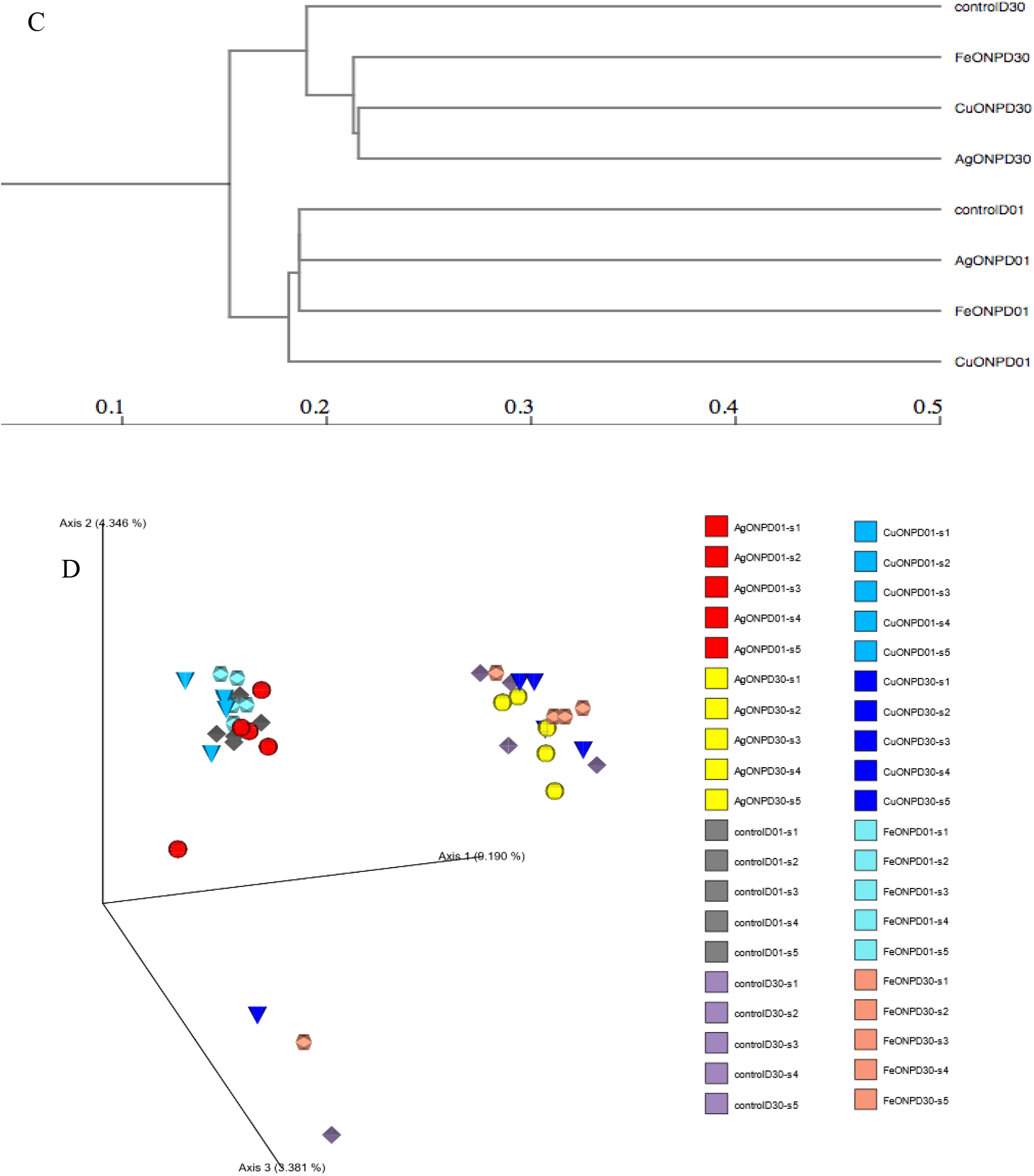
Principal coordinate analysis (PCoA) of soil bacteria calculated based on Bray-Curtis dissimilarity index (square-rooted to make metric) (a), Non-metric multi-dimensional scaling (NMDS) (b), showing bacteria diversity. Shapes represent different points of measurement while the colors represent the sample treatments. Hierarchical clustering dendrogram (c) of weighted UniFrac distance of bacterial communities showing the relationship between sample treatments treated with Cu_2_O, Fe_3_O_4_, and Ag_2_O NPs at 500 mg/kg soil on day **1** and day 30. (d) Bray-Curtis Emperor plot showing the clustering of samples along the first, second and third axes.

## Discussion

Soil bacteria are intricate part of soil processes such as nutrient recycling, decomposition of organic matter and improvement of plant growth^39^, therefore it is a major driver of ecosystem functioning. MONPs serve vital roles in industrial and consumer products and are known to have antimicrobial properties^10,11,40^. Given that some MONPs have the tendency to influence microbial activities and composition, this could pose potential risk to agroecosystems. With the sensitive nature of microbial diversity and structure to environmental stressors such as the presence of heavy metals and nutrient deficiency^14,41^, these factors become major assessment indicators of microbial response to environmental contaminant such as MONPs. This study provides results on the differential alteration of soil bacteria communities with MONPs. Using Cu_2_O, Fe_3_O_4_, and Ag_2_O NPs the impact of MONPs on soil microbial communities was evaluated.

The *Proteobacteria* phylum comprising of *alpha-* and *gamma-Proteobacteria* dominated the soil samples. The *alpha-Proteobacteria* were dominated by *Rhodospirillacease, Sphingomonadales, Rhizobiales*, bacteria orders while *gamma-Proteobacteria* were dominated by the *Xanthomonadaceae* and unclassified *gamma-Proteobacteria*. These bacteria groups known to be involved in nitrogen fixation^42^. These results are in consistent with other studies that observed dominance of *Proteobacteria* in agricultural soils^43,44^. Upon treatment with MONPs, there was no change in the abundance of *Proteobacteria* while *Bacteriodetes* generally increased after 30 days of exposure. The resistance of *Proteobacteria* to MONPs inhibitory effect could be that some classes of *Proteobacteria* are resistant to metal ions released from MONPs. For example, *Sphingomonadales* have been shown to have high tolerance to metal stress^45,46^. Ge et al^47^ reported a positive impact of TiO_2_ and ZnO NPs on *Sphingomonadaceae, Streptomycetaceae and Streptomyces* in dose-dependent manner. The stimulating effect of MONPs on some members of *Proteobacteria* such as *Sphingomonadales* could have overshadowed the reduction of few bacteria taxa thereby showing no change in abundance after 30 days of MONPs exposure. Because *alpha-Proteobacteria* are made up of soil nitrogen fixing bacteria and organic material decomposers, it is likely that the regulatory pathway for nitrogen fixation and carbon cycling were unaffected. *Bacteriodetes* generally increased after 30 days of soil incubation showing resilience in the presence of MONPs (Table 3). The increased abundance after 30 days could be a result of lack of penetration of MONPs into the bacteria cell. Similar stimulatory effect on *Bacteriodetes have been reported* which was attributed to the presence of metal ion resistance genes^27,48,49^. Studies have shown that the inhibitory effect of MONPs to gram-negative bacteria such as *E. coli* could be attributed to the structural make-up of the bacteria cell wall^50^. *Bacteriodetes* are reported to be a very important bacteria group responsible for degradation of organic materials such as chitin^49^, therefore their enrichment in the presence of MONPs supports the catabolic pathways and nutrient cycling.

Different classes of bacteria were stimulated by MONPs application for example the *Acdiobacteria-Gp10, Chloroflexi-unclassified, OD1_class_incertae_sedis* and *Flavobacteria* (Fig. 3). These results suggest that MONPs addition not only affect bacterial communities at lower levels of order, families, and genera as reported by Meli et al^51^, but also at the class and phylum level as shown in this study. Collins et al^52^ reported that control samples were dominated by members of *Rhizobiales, Flavobacteria* and *Sphingomonadales* but these bacteria groups decreased with the addition of copper and zinc oxide NPs and in soil depth over the time period. In contrast to our study, *Flavobacteria* was not detected in the control sample and on day 1 but only in MONPs treated samples at day 30. This shows variation in the differential composition of microbial communities that exist in agricultural soils, which could be dependent on the abiotic factors and agricultural practices such as fertilization, CO_2_ concentration, salinization and soil properties that shape the soil microbiome^53,54^.

In addition, the diversity measurement revealed increase in the bacterial community richness at day 1 with the addition of Fe_3_O_4_ NPs (Fig. 5a) compared to control. The presence of Fe_3_O_4_ NPs in the soil stimulates the growth of some bacterial groups such as *beta-Proteobacteria* as shown using qPCR method (Fig. 1c). Iron oxide NPs have surface sites that binds to soil organic compounds such as humic and fluvic acids that are intricate part of the soil microbiome, therefore could promote the bioavailability of iron to soil bacteria thus stimulating the growth of soil microbes^55^. After 30 days of MONPs exposure, alpha diversity measured by Chao1 index revealed that the bacteria richness reduced with the addition of MONPs and was greater with Ag_2_O and Cu_2_O NPs (Fig. 5). The length of MONPs contributed to the degree of MONPs impact on bacteria community diversity and composition (Table 4). Given that the Ag_2_O and Cu_2_O NPs used in this study are known to have antimicrobial properties^11,40^, we proposed that the reduction effects on the bacterial diversity was a result of the release of metal ions from MONPs thereby causing bacterial toxicity.

In consistent with our results, other studies have reported a shift in the relative abundance of different bacteria phyla after 30 days of Ag, CuO and ZnO NPs exposure compared to 2 h of NP exposure^41^. Accordingly, You et al^56^ monitored a similar significant decrease in bacteria richness after 30 days of 0.5, 1 and 1 mg/g soil ZnO NPs exposure. Another study observed a greater decrease on soil bacterial communities at day 60 than day 15 in TiO_2_ and ZnO NPs treated soils but had no effect at day zero^22^. The PC_o_A and NMDS at the OTU level (97% similarity) showed distinct difference in bacterial community composition at day 1 and day 30 confirmed by AMOVA analysis (p < 0.001). Close bacterial clustering was observed on day 1 of MONPs exposure indicating that the bacterial communities were stable (Fig. 6). After 30 days of exposure, MONP treated soil samples were significantly distant from control sample indicating a significant change in the bacterial community. Overall, the application of MONPs caused a substantial shift in bacteria community composition and structure which was driven by the duration of MONPs exposure and the type of MONPs.

In conclusion, changes in the relative abundance of known bacterial taxa and overall bacterial diversity with Cu_2_O, Fe_3_O_4_ and Ag_2_O NPs at 500 mg/kg soil over a 30-day period of exposure was investigated using 16S rRNA sequencing and qPCR techniques. Our results showed that the addition of MONPs altered bacterial community composition by causing significant reduction in bacterial diversity and change in bacterial abundance. After 30 days of MONPs exposure, the bacterial diversity was significantly reduced with Cu_2_O and Ag_2_O NPs having greater impact. The *alpha-* and *beta-Proteobacteria* and *Bacteriodetes* were not negatively impacted in the presence of MONPs over the 30-day period of exposure thus, supports the nitrogen fixation and chitin degradation pathway. The long exposure of microbial communities to MONPs contributed to the degree of bacteria diversity reduction effects observed in this study. The variation in susceptibility and sensitivity of some bacteria groups to MONPs stimulated a shift in microbial community structure towards a more MONPs tolerant bacteriome (e.g *Bacteriodetes)*. Since soil serve as a sink to NPs deposition, more long-term studies of NPs exposure to soil microbiome is required to determine the concentration threshold that could substantially influence key bacteriome and the functioning of agroecosystems.

